## Supplementary figures and images for "HSV-1 and influenza infection induce linear and circular splicing of the long NEAT1 isoform"

### S1 Fig

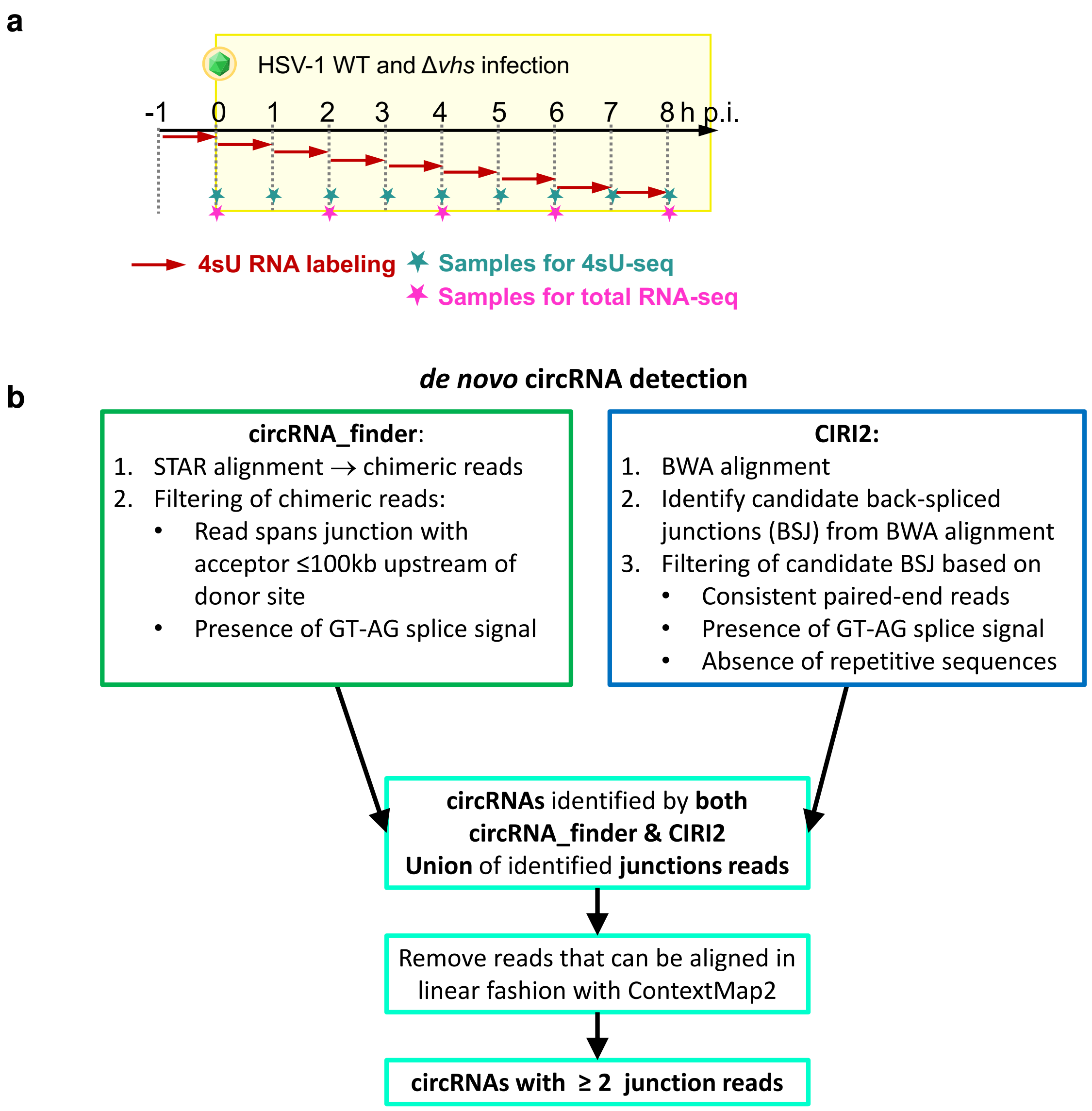

### S2 Fig

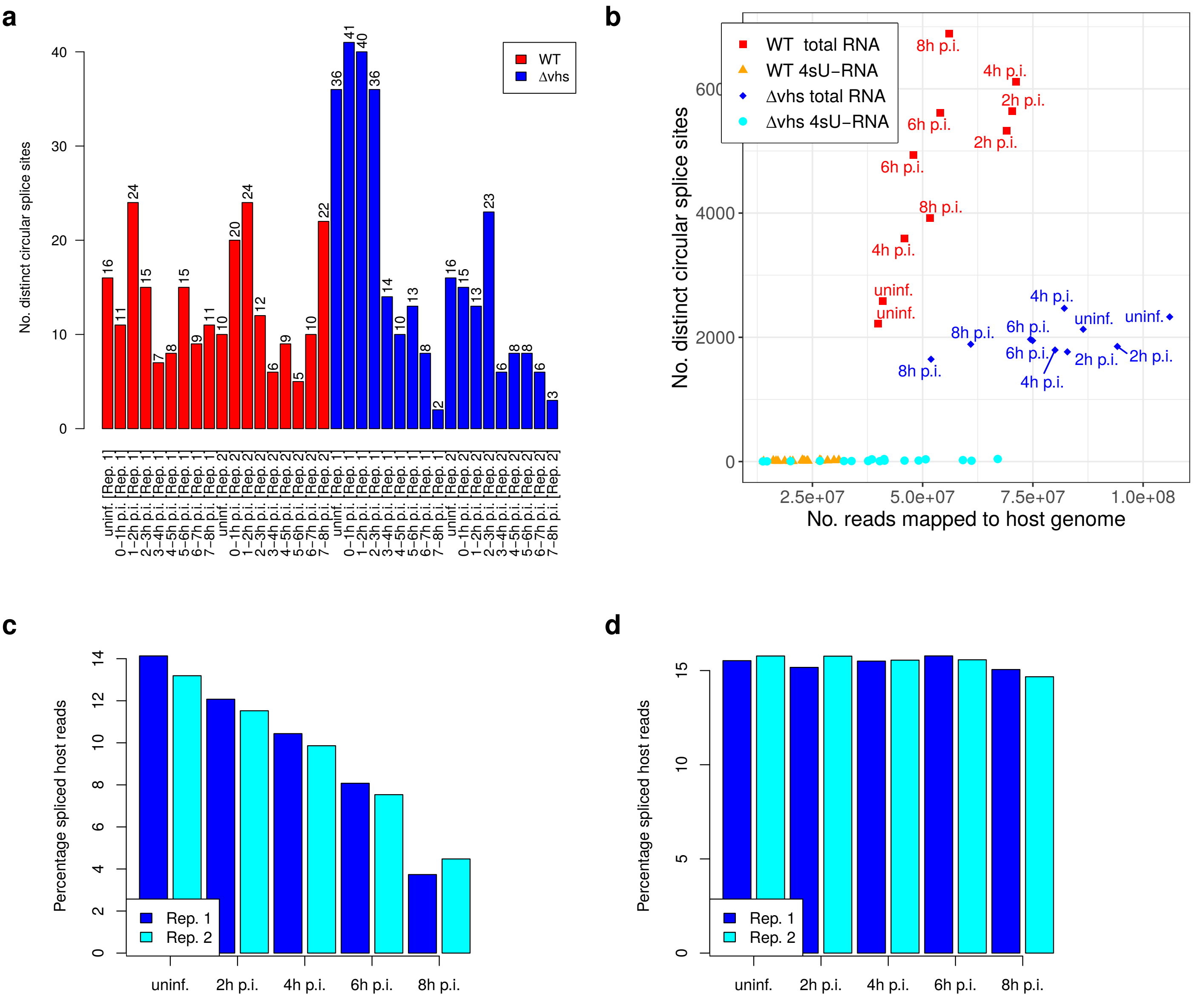

### S3 Fig

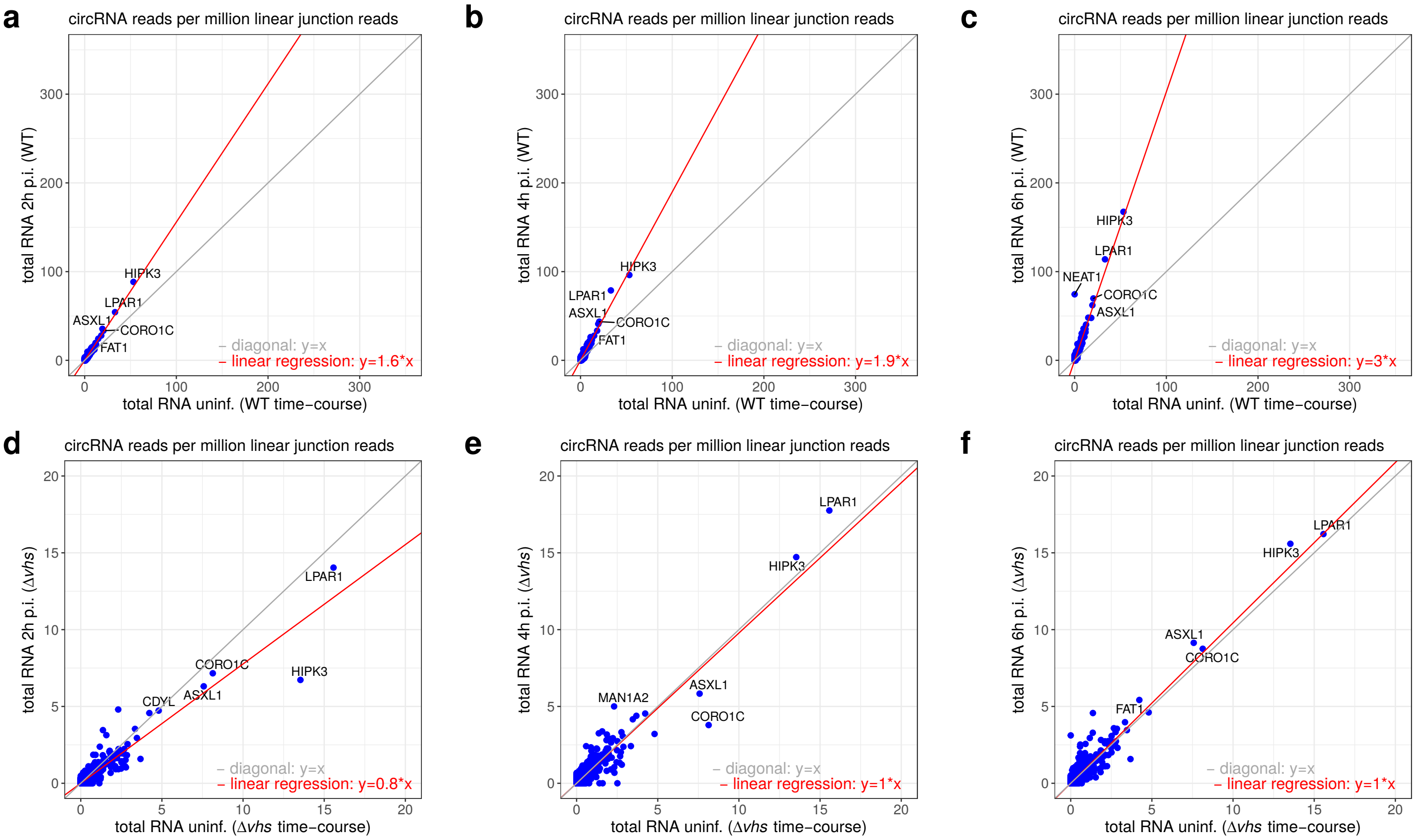

### S4 Fig

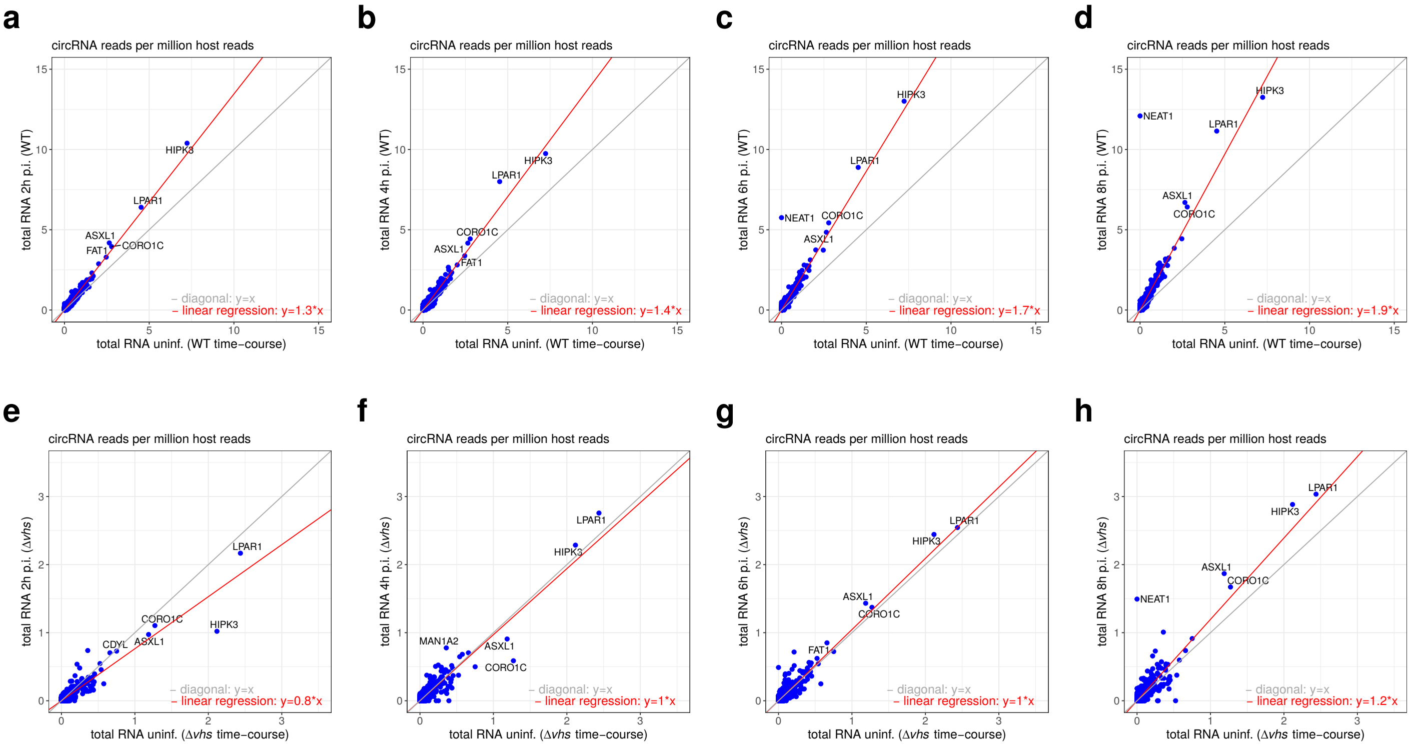

### S5 Fig

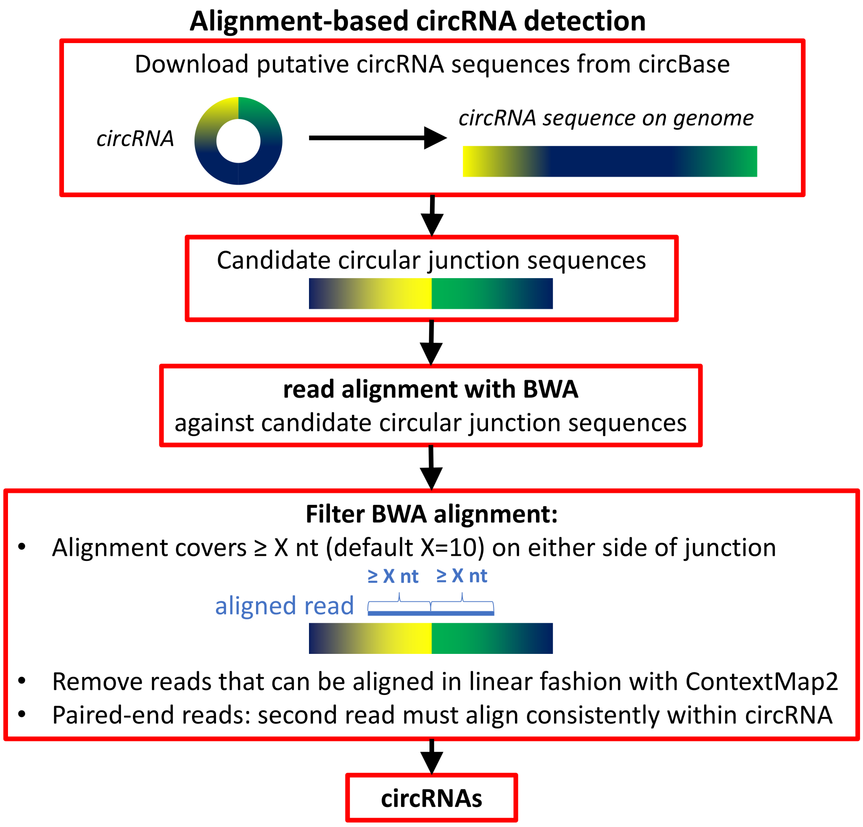

### S6 Fig

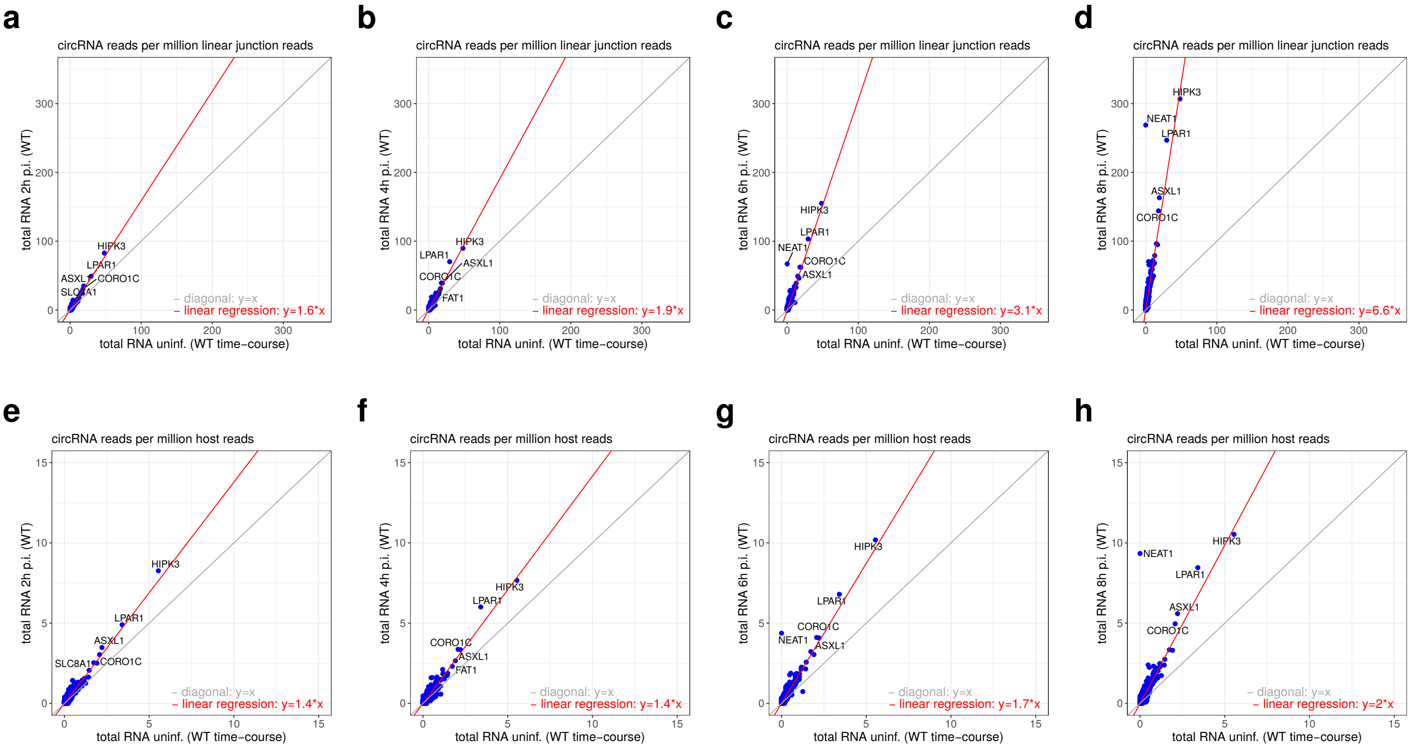

### S7 Fig

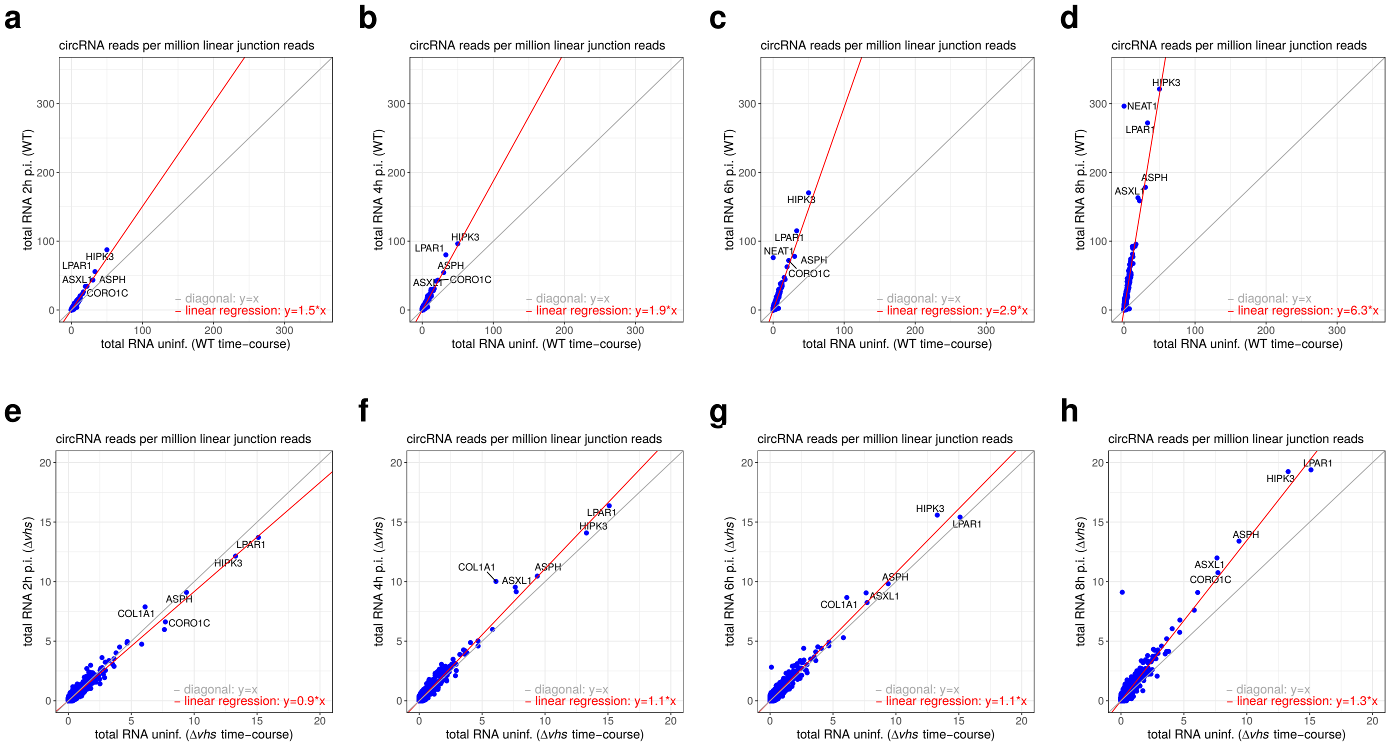

### S8 Fig

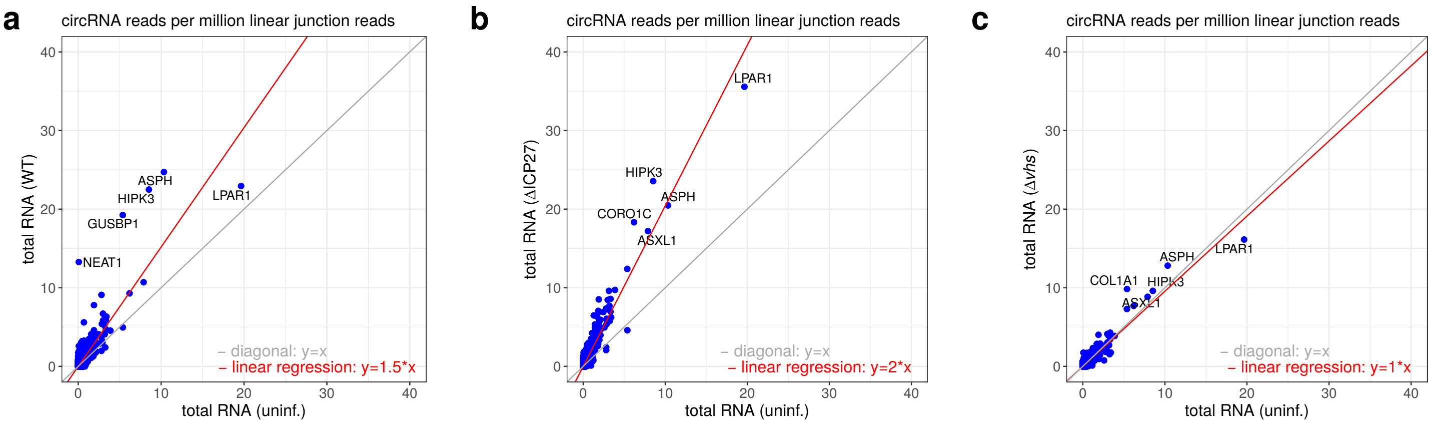

### S9 Fig

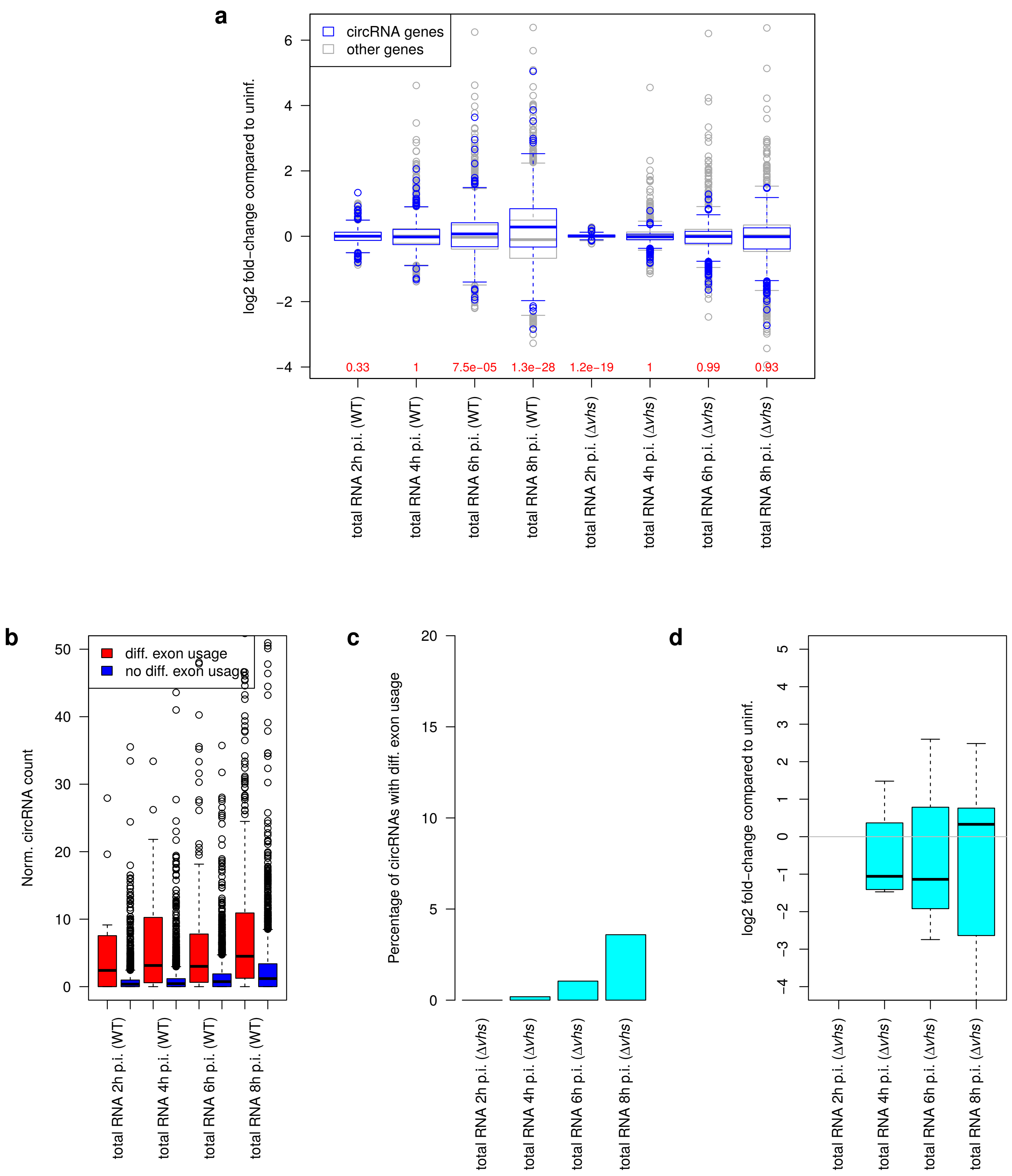

### S10 Fig

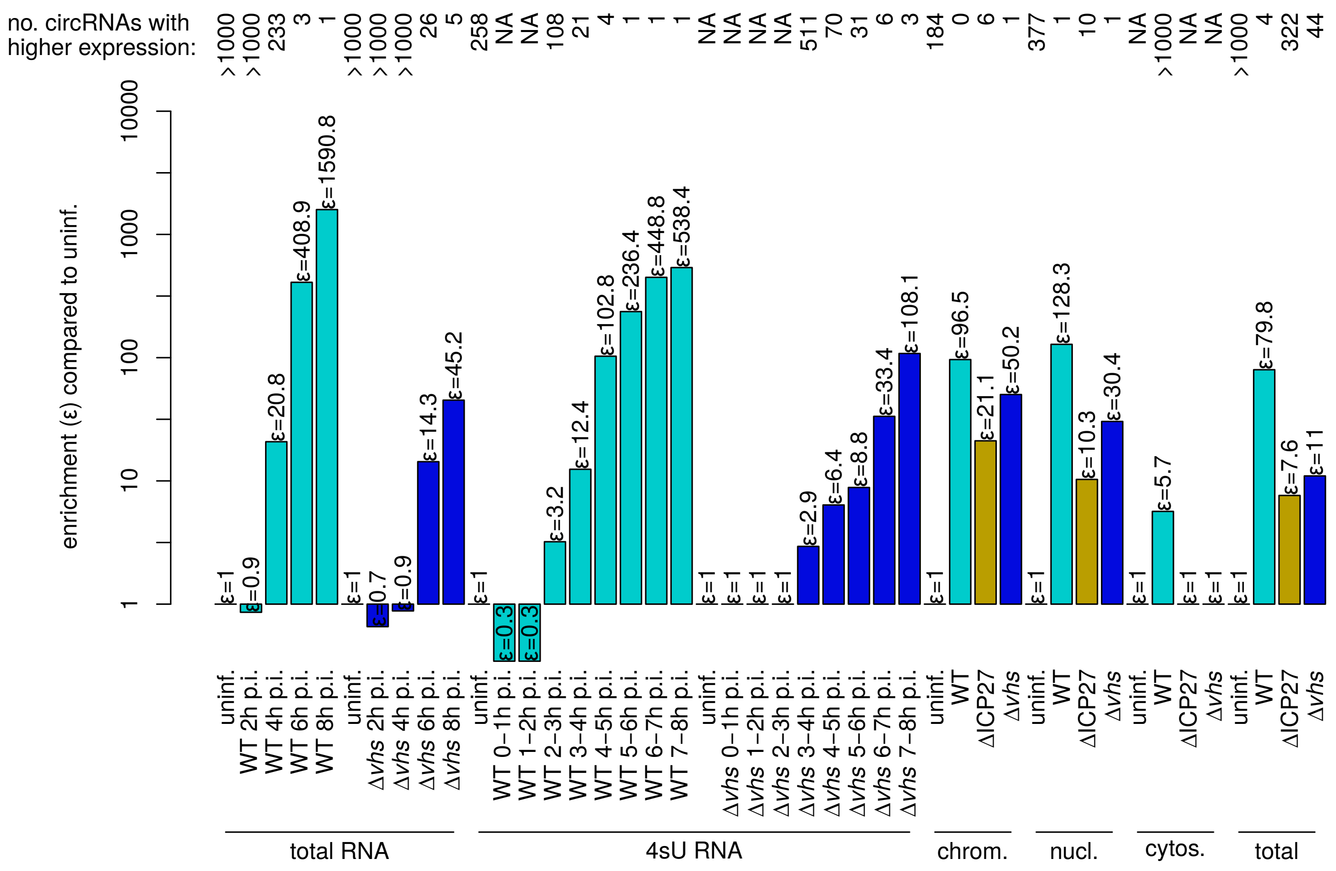

### S11 Fig

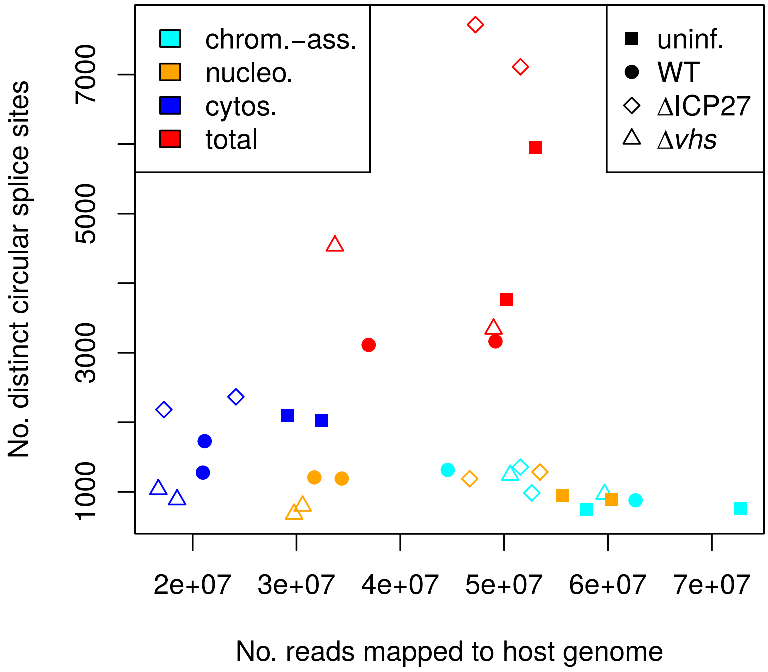

### S12 Fig

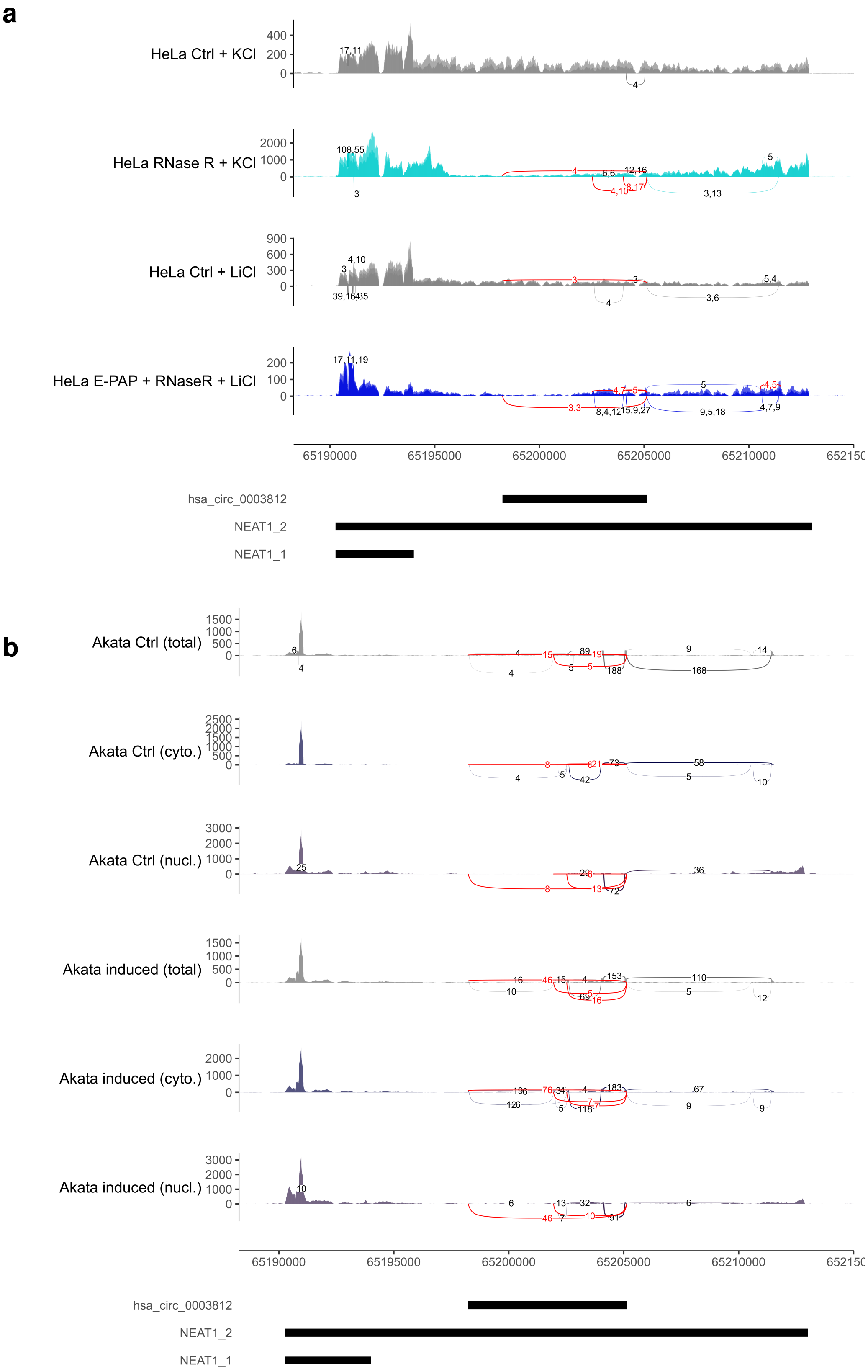

### S13 Fig

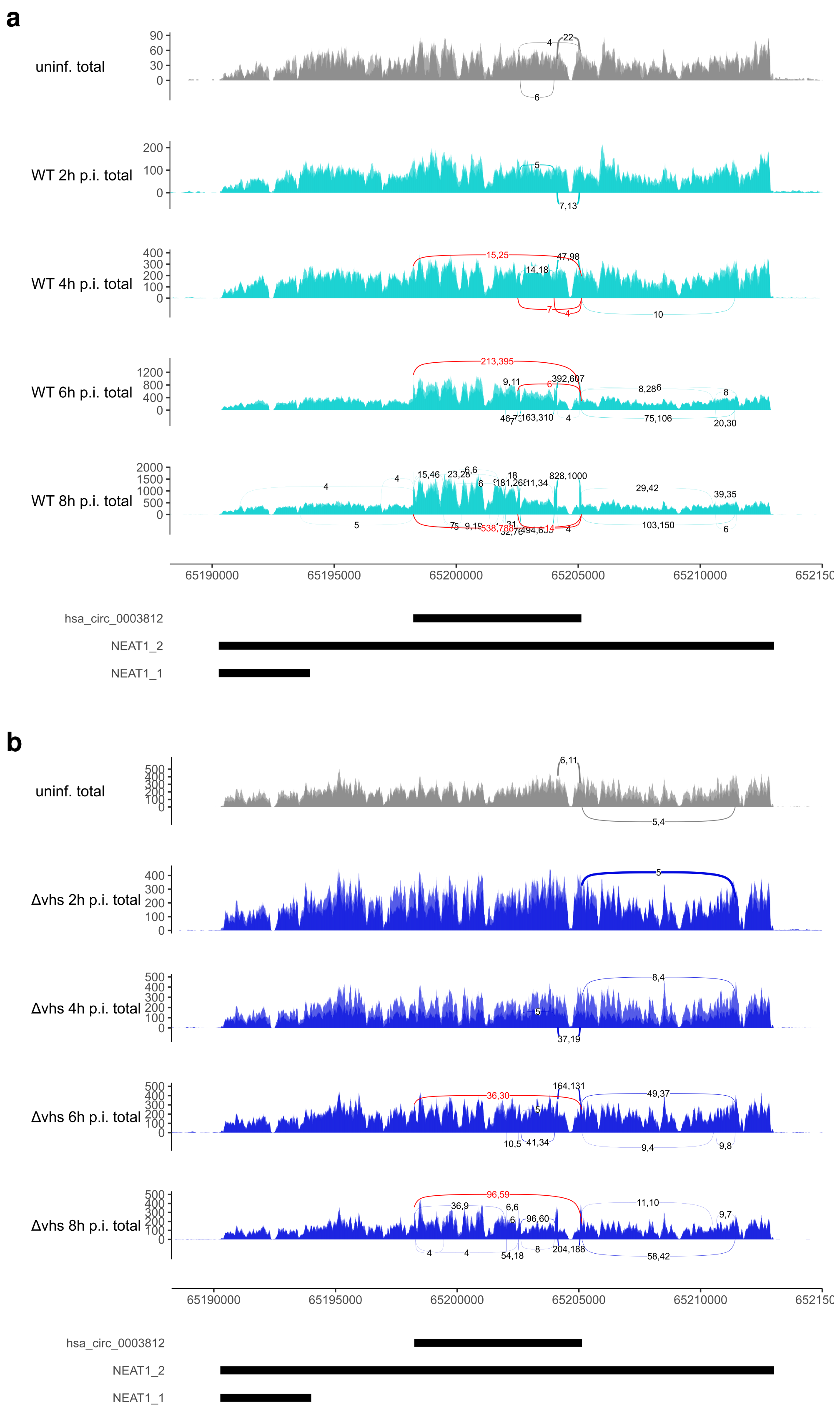

### S15 Fig

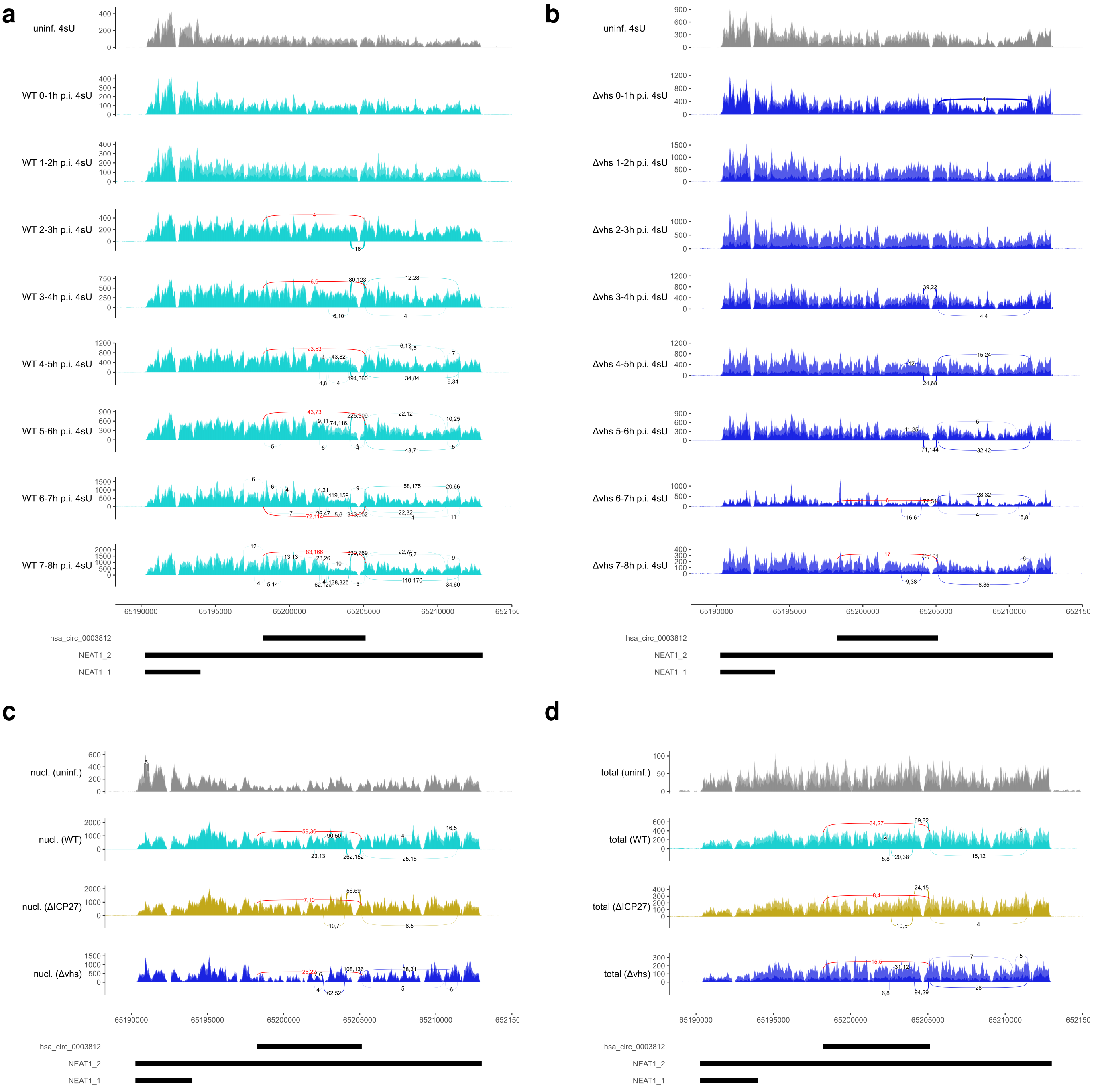

### S16 Fig

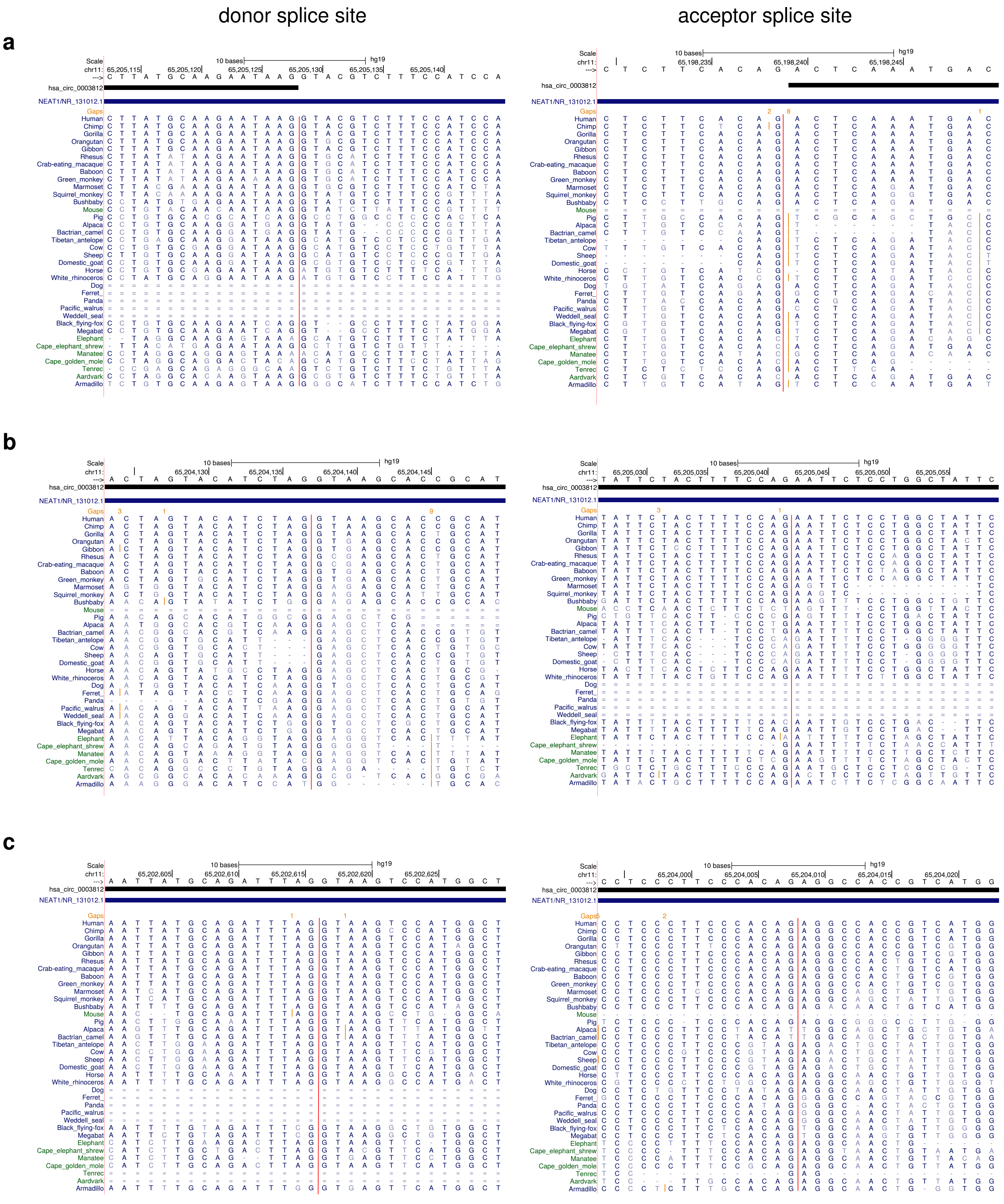

### S17 Fig

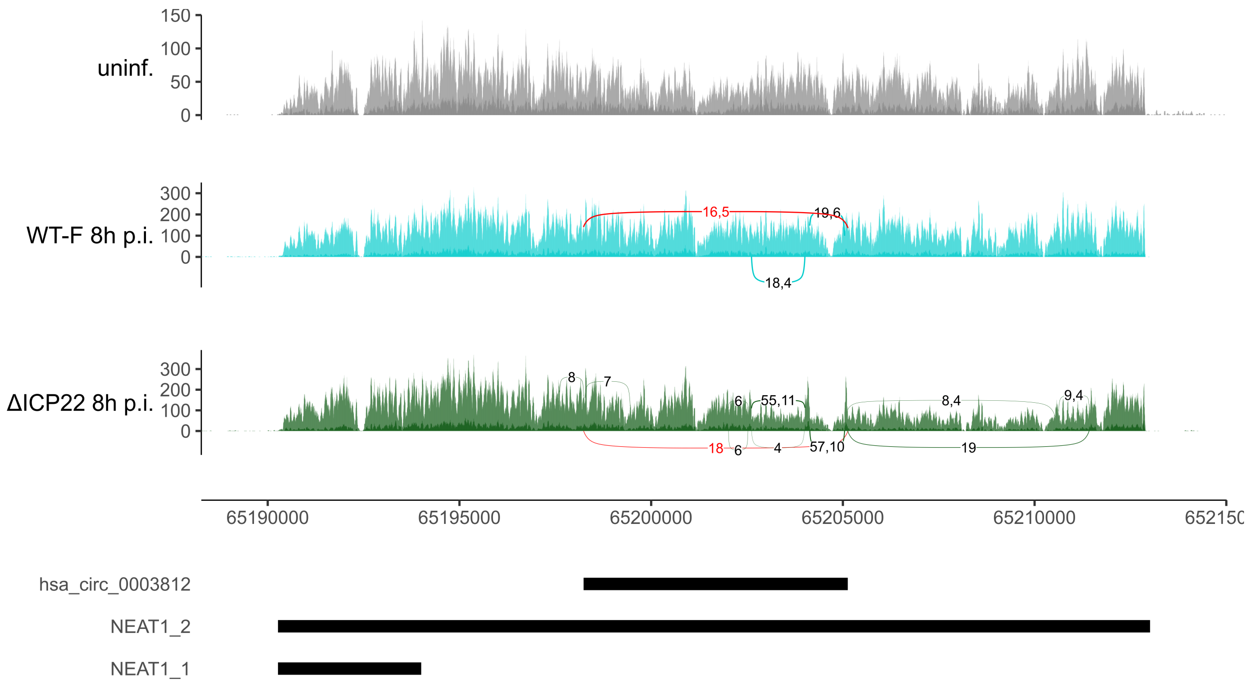

### S18 Fig

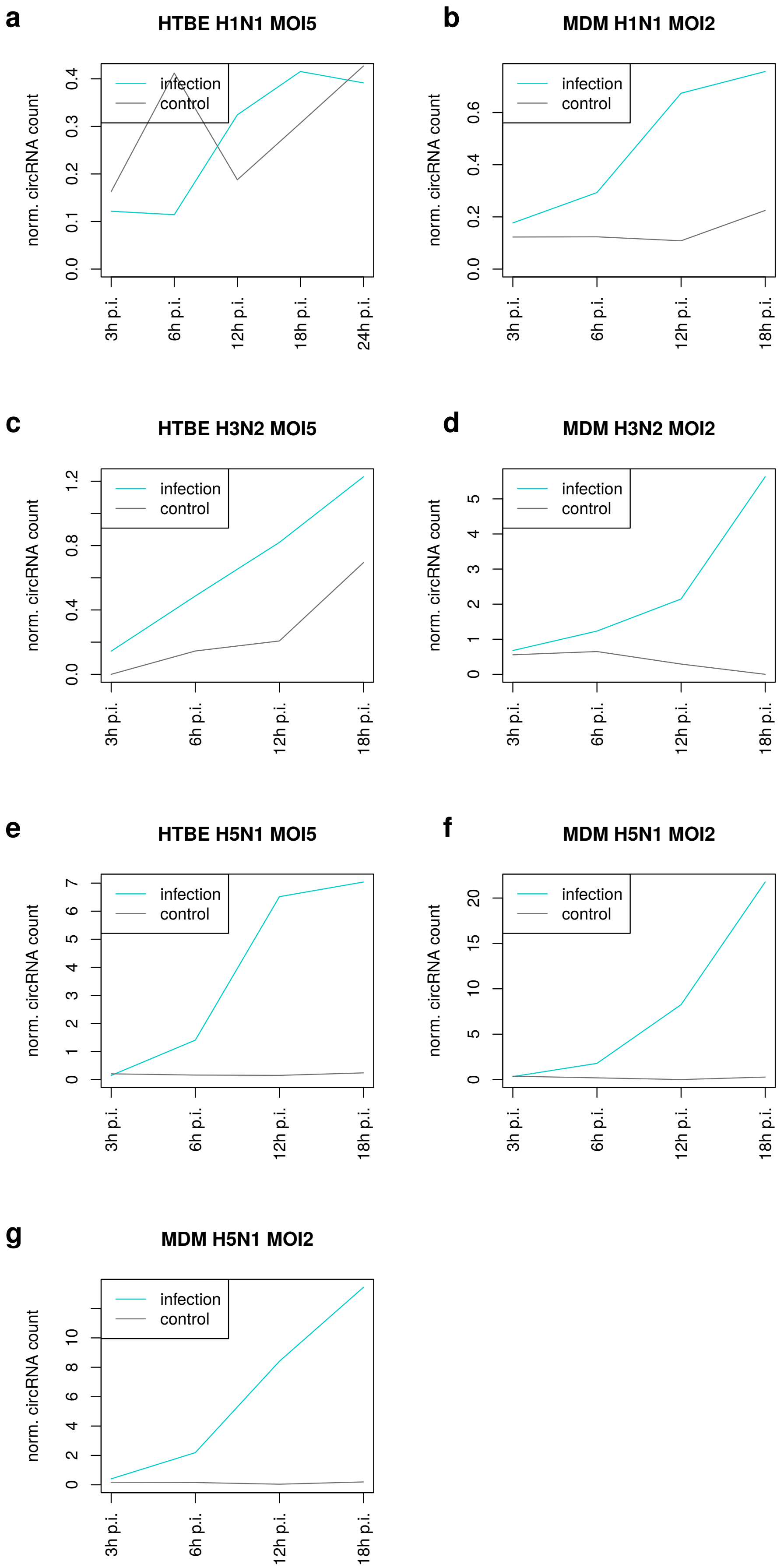

### S19 Fig

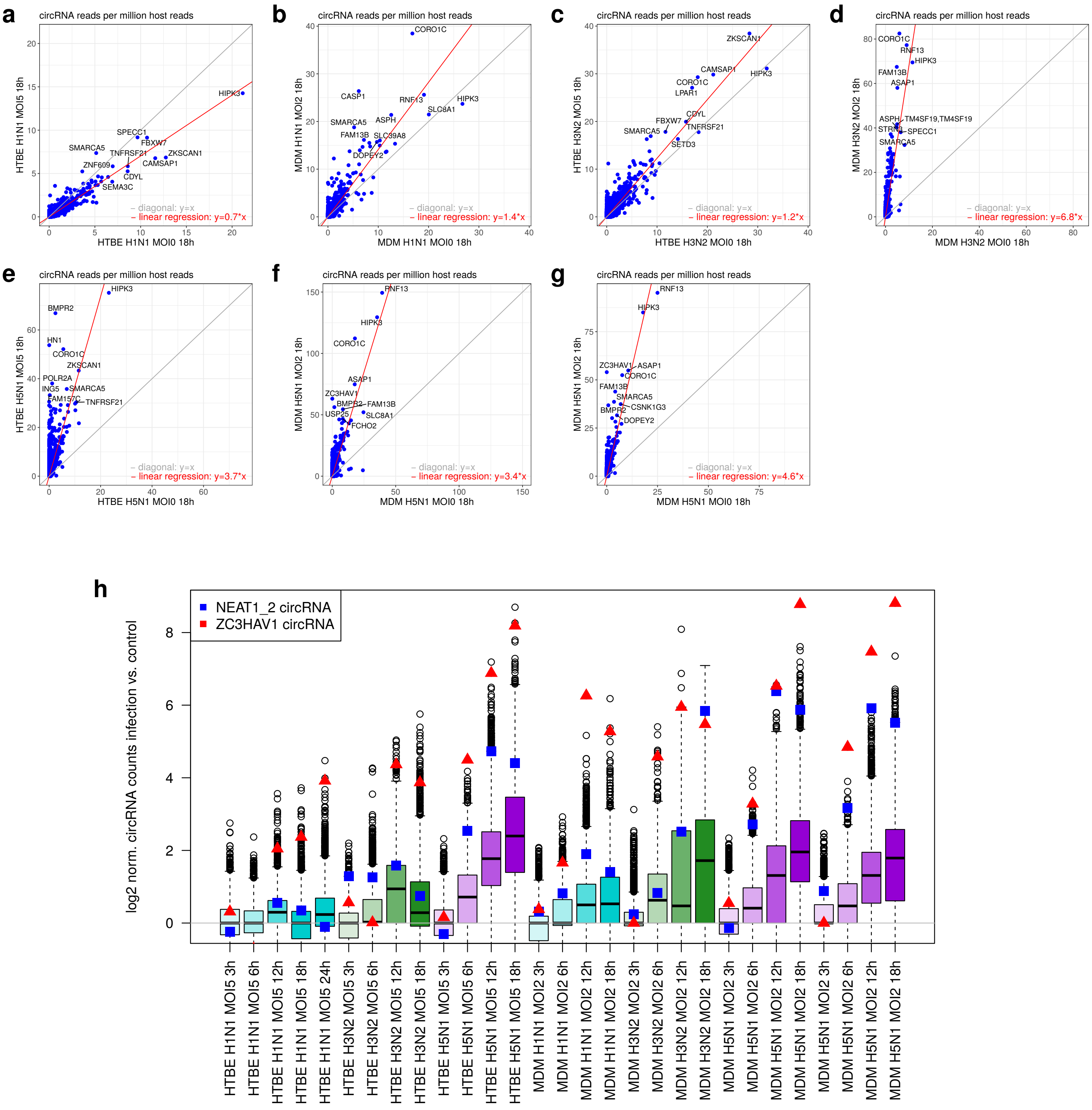

### S20 Fig

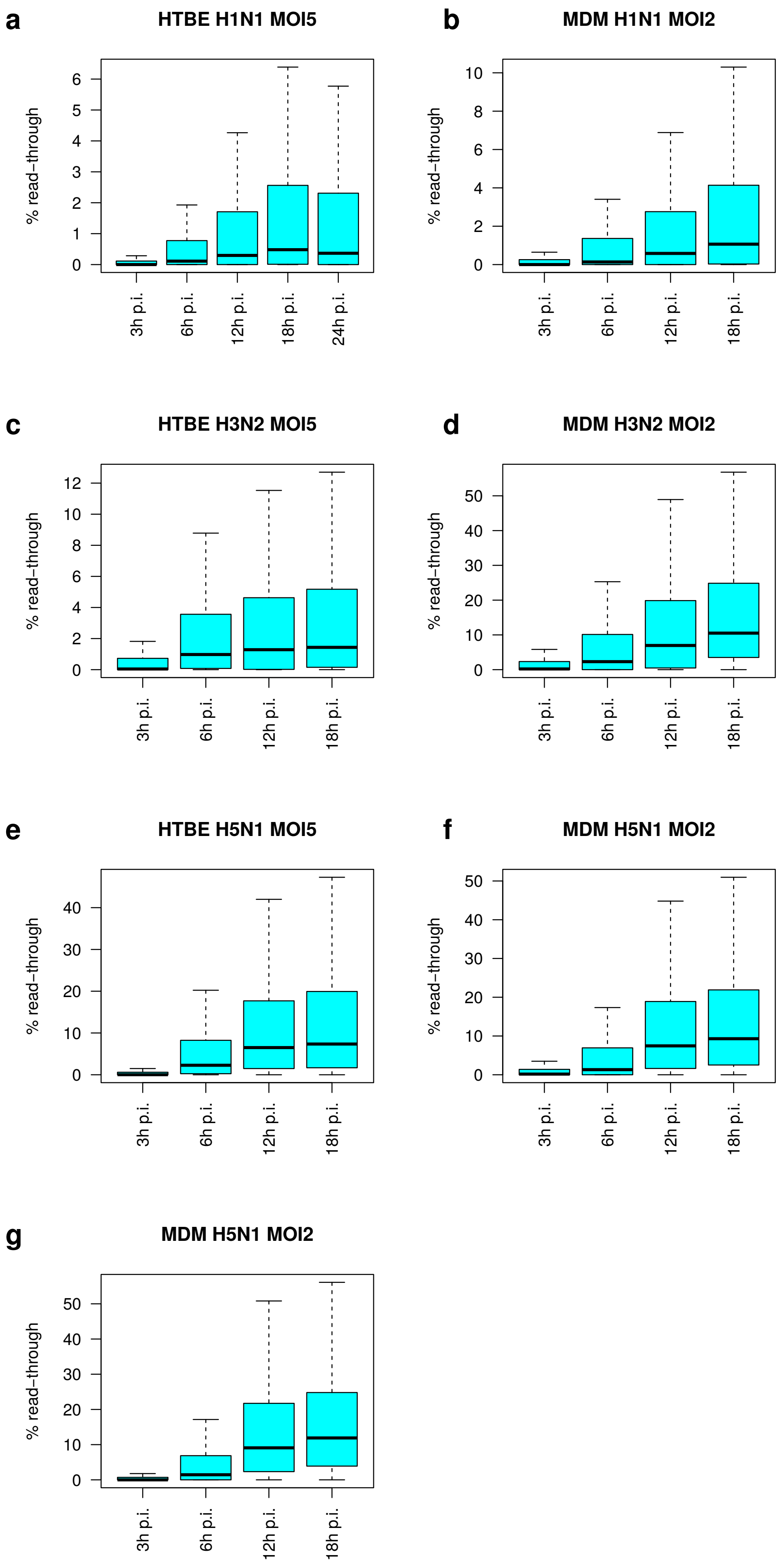

### S22 Fig

a

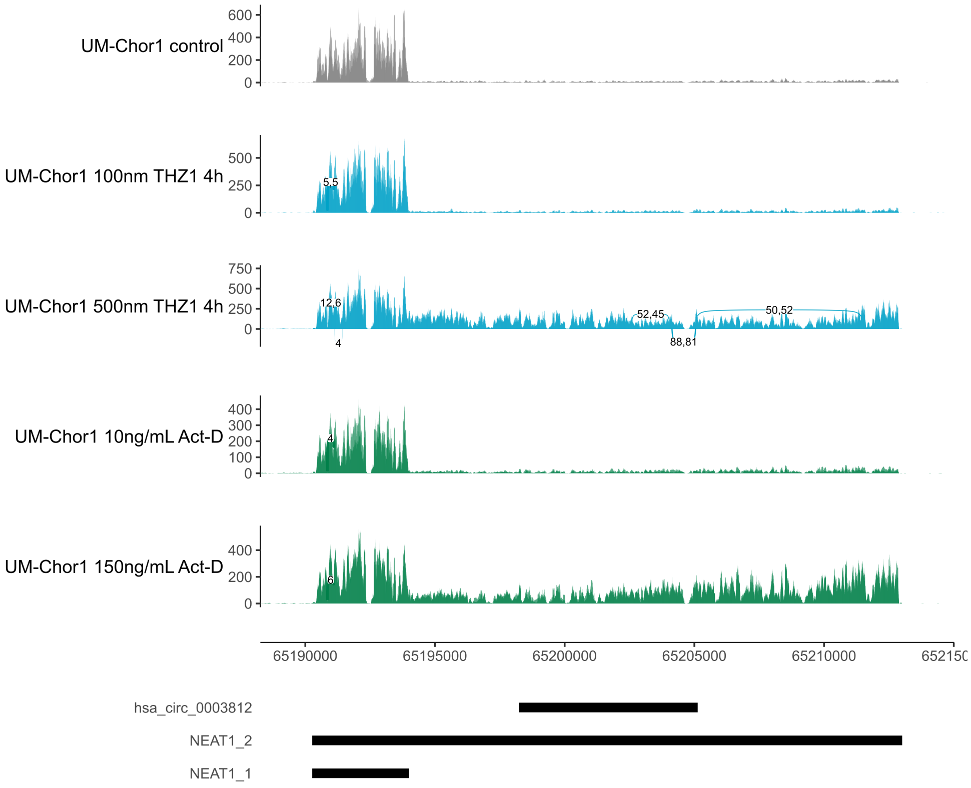


b

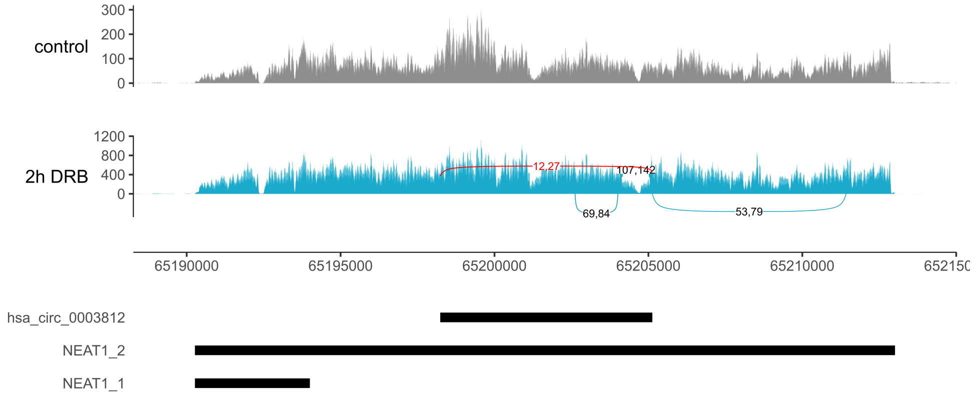


#### S22 Figure

### S23 Fig

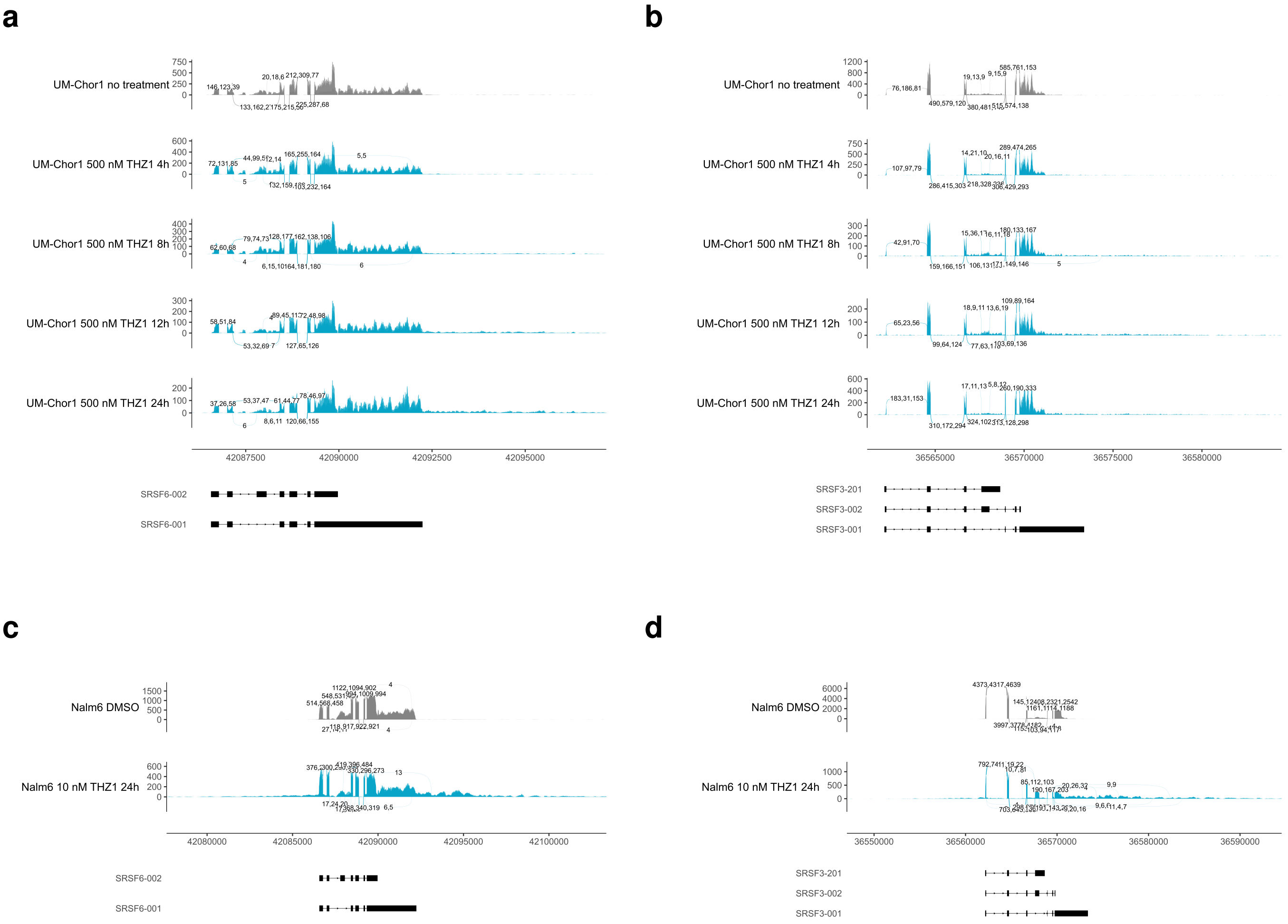
