## Supplementary material for "HSV-1 and influenza infection induce linear and circular splicing of the long NEAT1 isoform": S14 Fig

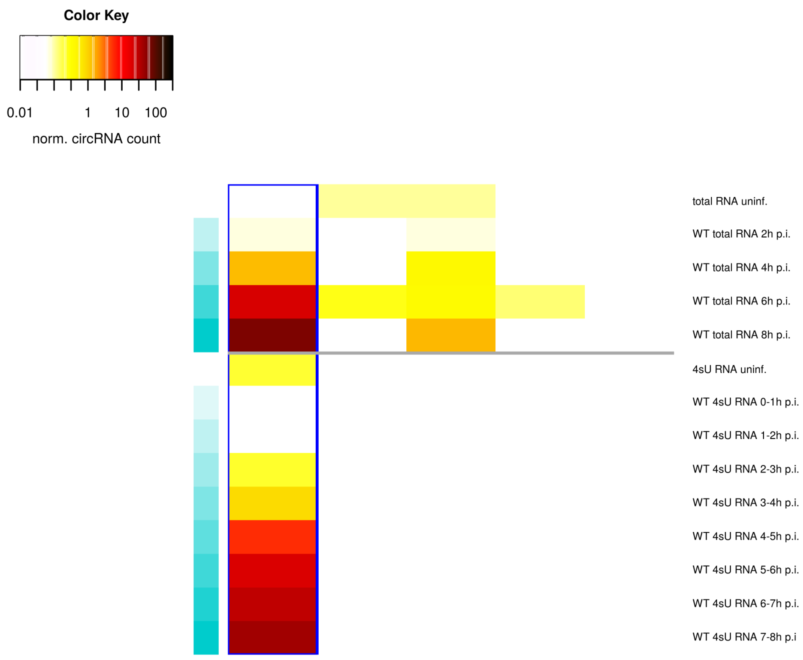


**b**

| | no. circular reads (40 nt ov. on both sides) | no. confirming read pairs |
|---|---|---|
| **WT total RNA** | | |
| uninf. (1) | 0 | 0 |
| uninf. (2) | 0 | 0 |
| 2h p.i. (1) | 1 | 0 |
| 2h p.i. (2) | 0 | 0 |
| 4h p.i. (1) | 5 | 0 |
| 4h p.i. (2) | 9 | 2 |
| 6h p.i. (1) | 54 | 7 |
| 6h p.i. (2) | 121 | 22 |
| 8h p.i. (1) | 120 | 11 |
| 8h p.i. (2) | 170 | 45 |
| **WT 4sU RNA** | | |
| uninf. (1) | 0 | 0 |
| uninf. (2) | 1 | 0 |
| 0-1h p.i. (1) | 0 | 0 |
| 0-1h p.i. (2) | 0 | 0 |
| 1-2h p.i. (1) | 0 | 0 |
| 1-2h p.i. (2) | 0 | 0 |
| 2-3h p.i. (1) | 0 | 0 |
| 2-3h p.i. (2) | 1 | 0 |
| 3-4h p.i. (1) | 2 | 0 |
| 3-4h p.i. (2) | 1 | 0 |
| 4-5h p.i. (1) | 5 | 0 |
| 4-5h p.i. (2) | 9 | 0 |
| 5-6h p.i. (1) | 12 | 2 |
| 5-6h p.i. (2) | 22 | 0 |
| 6-7h p.i. (1) | 18 | 1 |
| 6-7h p.i. (2) | 24 | 2 |
| 7-8h p.i. (1) | 23 | 1 |
| 7-8h p.i. (2) | 44 | 8 |

**c**

**Identification of confirming read pairs**

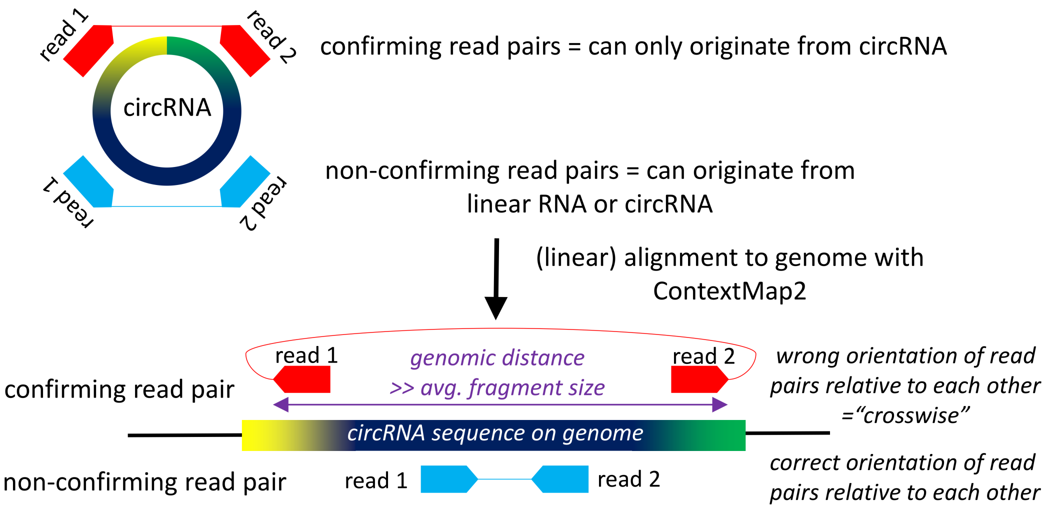


**S14 Figure**
