## Supplementary material for "HSV-1 and influenza infection induce linear and circular splicing of the long NEAT1 isoform": S21 Fig

**a**

| | no. projects | no. samples |
|---|---|---|
| **all results** | 909 | 6391 |
| **pattern in description** | | |
| (HSV-1\|herpes simplex virus) | 7 | 36 |
| (influenza\|H1N1\|H3N2\|H5N1) | 7 | 63 |
| (circRNA\|circular RNA) | 62 | 344 |
| RNase R | 21 | 136 |
| blood | 179 | 2417 |
| platelet | 13 | 65 |
| erythrocyt | 3 | 27 |
| monocyt | 60 | 541 |
| leu[ck]ocyt | 13 | 85 |
| lymphocyt | 22 | 137 |
| macrophag | 42 | 333 |
| leukem | 53 | 356 |
| PBMC | 34 | 323 |
| K562 | 21 | 497 |
| (CDK7\|THZ1) | 9 | 26 |

**b**

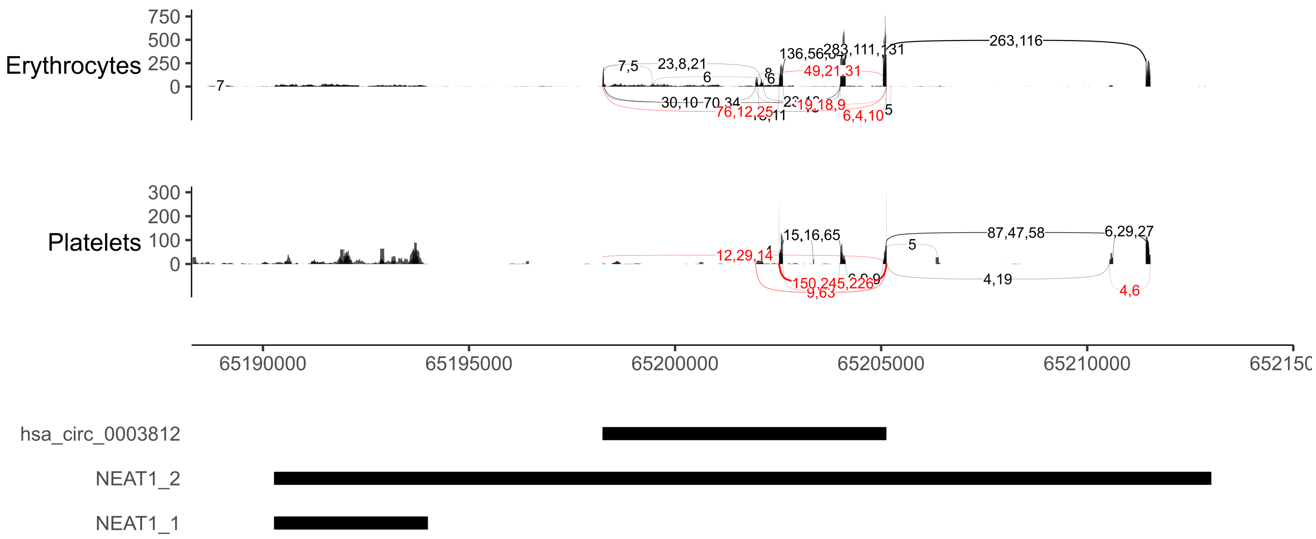


**S21 Figure**
